# Apparent Motion Perception in the Praying Mantis: Psychophysics and Modelling

**DOI:** 10.1101/320606

**Authors:** Ghaith Tarawneh, Lisa Jones, Vivek Nityananda, Ronny Rosner, Claire Rind, Jenny Read

**Affiliations:** Institute of Neuroscience, Newcastle University

## Abstract

Apparent motion is the perception of a motion created by rapidly presenting still frames in which objects are displaced in space. Observers can reliably discriminate the direction of apparent motion when inter-frame object displacement is below a certain limit, Dmax. Earlier studies of motion perception in humans found that Dmax scales with spatial element size, interpreting the relationship between the two as linear, and that Dmax appears to be lower-bounded at around 15 arcmin. Here, we run corresponding experiments in the praying mantis *Sphodromantis lineola* to investigate how Dmax scales with element size. We used moving random chequerboard patterns of varying element and displacement step sizes to elicit the optomotor response, a postural stabilization mechanism that causes mantids to lean in the direction of large-field motion. Subsequently, we calculated Dmax as the displacement step size corresponding to a 50% probability of detecting an optomotor response in the same direction as the stimulus. Our main findings are that mantis Dmax appears to scale as a power-law of element size and that, in contrast to humans, it does not appear to be lower-bounded. We present two models to explain these observations: a simple high-level model based on motion energy in the Fourier domain and a more detailed one based on the Reichardt Detector. The models present complementary intuitive and physiologically-realistic accounts of how Dmax scales with element size in insects.

**Author Summary:** Computer monitors, smart phone screens and other forms of digital displays present a series of still images (frames) in which objects are displaced in small steps, tricking us into perceiving smooth motion. This illusion is referred to as “apparent motion”, and for it to work effectively the magnitude of each displacement step must be smaller than a certain limit, referred to as Dmax. Previous studies have investigated the relationship between this limit and object size in humans and found that larger objects can be displaced in larger steps without affecting motion perception. In this work, we investigated the same relationship in the praying mantis *Sphodromantis lineola* by presenting them with moving chequerboard patterns on a computer monitor. Even though motion perception in humans and insects are believed to be explained equally well by the same underlying model, we found that Dmax scales with object size differently in mantids. These results suggest that there may be qualitative differences in how mantids perceive apparent motion compared to humans.

## 1 Introduction

Detecting motion is an important visual function in all animals. Movement can signal the presence of prey, predator or a conspecific in the animal’s surrounding environment and is therefore an important cue for many forms of behavior. To understand how motion detection in animal visual systems work it is helpful to first examine and understand their responses to basic forms of visual input such as apparent motion stimuli. Apparent motion is an illusion of motion created by the rapid presentation of successive still frames in which objects are gradually displaced in space. The ability of an observer to perceive motion in such stimuli depends on a number of factors including the frame rate of presentation and the amount of spatial displacement between individual frames. When inter-frame displacement is made large, for example, objects no longer appear as smoothly moving and are instead perceived to be jumping in discrete steps. Increasing inter-frame displacement further, beyond a certain limit, makes observers unable to tell motion direction. This limit is known as Dmax (Braddick, 1974) and is a characteristic feature of a visual motion detection system. Measuring Dmax and examining its relationship with other stimuli features can reveal details about the computational mechanisms of motion detection in the visual system. It can also improve our understanding of how apparent motion is perceived and offer ways to improve video display technologies such as TVs and computer monitors that rely on this phenomenon.

In an early study of apparent motion perception in humans, subjects were presented with random dot patterns and tested on their ability to identify a correlated region of dots that was uniformly displaced on each frame (Braddick, 1974). The individual stimulus frames contained no cues to the region’s location and so subjects could only distinguish it based on the motion of its constituent dots. The study found that the subjects could not detect motion when the region was displaced by more than 15 arcmins between frames, and that varying dot size in the range 2 to 11 arcmins had no effect on this result. The author postulated that humans have a constant Dmax of 15 arcmins that is imposed by the dimensions of elementary motion detectors in the human visual system. Subsequent studies however have shown that, while Dmax appears to be constant for element sizes up to 15 arcmins (consistent with (Braddick, 1974)), increasing element size further caused a proportional increase in Dmax (Cavanagh et al., 1985; Morgan, 1992). Absolute and relative Dmax values were presumed to be in favor of different explanations of how humans perceive motion (Braddick, 1974; Chang and Julesz, 1983; Sato, 1989) but were later shown to be accounted for by a simple model consisting of an initial spatial filtration stage followed by a motion detection mechanism dependent on element size (Morgan, 1992). Although many studies have investigated Dmax and its implications in humans over the course of four decades (Braddick, 1974; Morgan, 1992; Tripathy et al., 2012), no corresponding studies have been reported in insects. Measuring the properties of Dmax would not only give an insight into the operation of visual motion detection in insects but would also enable comparison with humans, potentially shedding light on the evolutionary origins of motion perception.

In this study, we tested apparent motion perception in the praying mantis *Sphodromantis lineola* using moving random chequerboard patterns rendered on a computer monitor. We measured Dmax for a range of element sizes and found that insect Dmax scales with element size. This scaling is observed in humans for large sizes, but whereas human Dmax asymptotes to a fixed minimum at fine scales, insect Dmax continued to decrease as element size decreased, down to the smallest scales detectable by the insects.

We present two computational models that account for the relationship between Dmax and element size in the mantis. The first is based on the correlation-type or Reichardt Detector (RD) that is widely used to account for large field motion perception in insects (Borst, 2014a; Hassenstein and Reichardt, 1956) and is in good agreement with our observations both qualitatively and quantitatively. The second is a simplified energy-based model (Adelson and Bergen, 1985) that, although not assuming limited spatial and temporal sensitivity similar to the RD, still accounts for the main qualitative features of our results. Taken together, the models offer complementary insights into insect Dmax: the first shows that experimental observations are consistent with biophysically-realistic models of motion detection while the second offers corresponding intuitive explanations based on motion energy distribution in the Fourier spectrum.

## 2 Results

### 2.1 Experimental Findings

Mantids were placed in front of a Cathode Ray Tube (CRT) monitor and viewed a full screen random chequer-board stimulus moving horizontally in a series of steps. The size of the individual chequers (the *element size*) and the displacement magnitude in each step (the *step size*) were varied in different trials (Figure 1). When the step size was small, the stimulus appeared to humans as a smoothly moving pattern and elicited a postural stabilization mechanism (the optomotor response) causing mantids to lean in the direction of motion. When the step size was made larger however the stimulus appeared to humans as a still image moving in discrete steps and mantids responded by either peering (moving the head and body from side to side, simultaneously counter-rotating the head so as to keep the eyes looking straight) or remaining still. In all trials, the time between steps was proportional to step size so apparent motion speed was kept constant at 12.5 cm/s. Figure 2 shows stimulus space time plots and corresponding spatiotemporal Fourier spectra for different step sizes.

**Figure 1:**
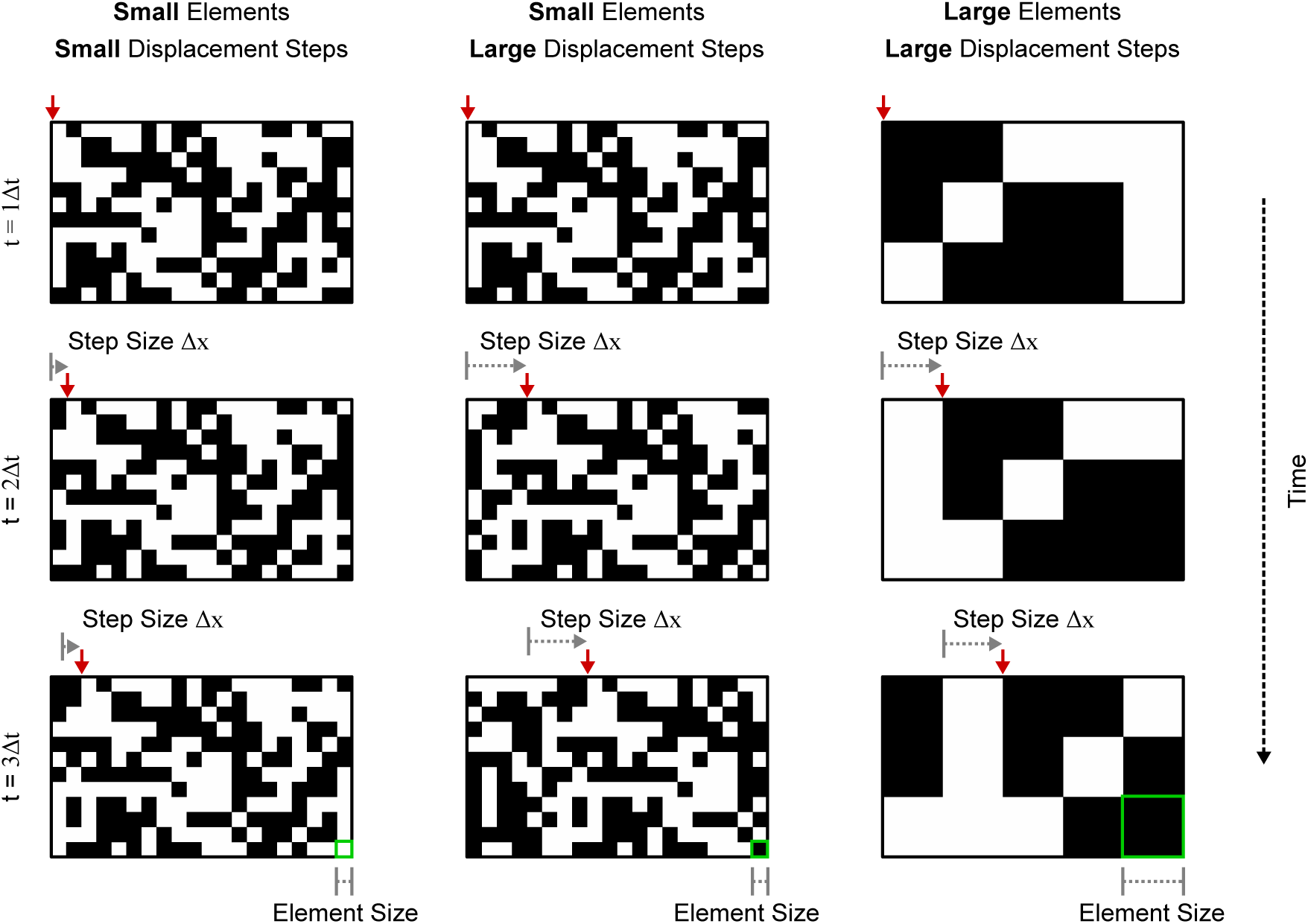
Examples of the random chequerboard pattern stimuli used in the experiment. Each column of panels shows still frames of a random chequerboard pattern that moves across a screen. The pattern is displaced by a fixed step Δ*x* on Δ*t* intervals where the speed *v* = ∆*x*/Δ*t* is constant (the red arrow at the top of each panel points to a reference point for easier visualization). The perception of motion in these patterns is strongly dependent on the displacement step Δ*x* relative to the pattern’s chequer/element size. When Δ*x* is comparable to element size, humans perceive the pattern as moving smoothly and can identify its direction reliably (column 1). Increasing Δ*x* causes pattern motion to appear more “jerky” and makes its direction harder to identify (column 2). Because the perception of motion is dependent on the ratio between element and step size, increasing element size to match step size makes the patterns appear to be moving smoothly again (column 3).

**Figure 2:**
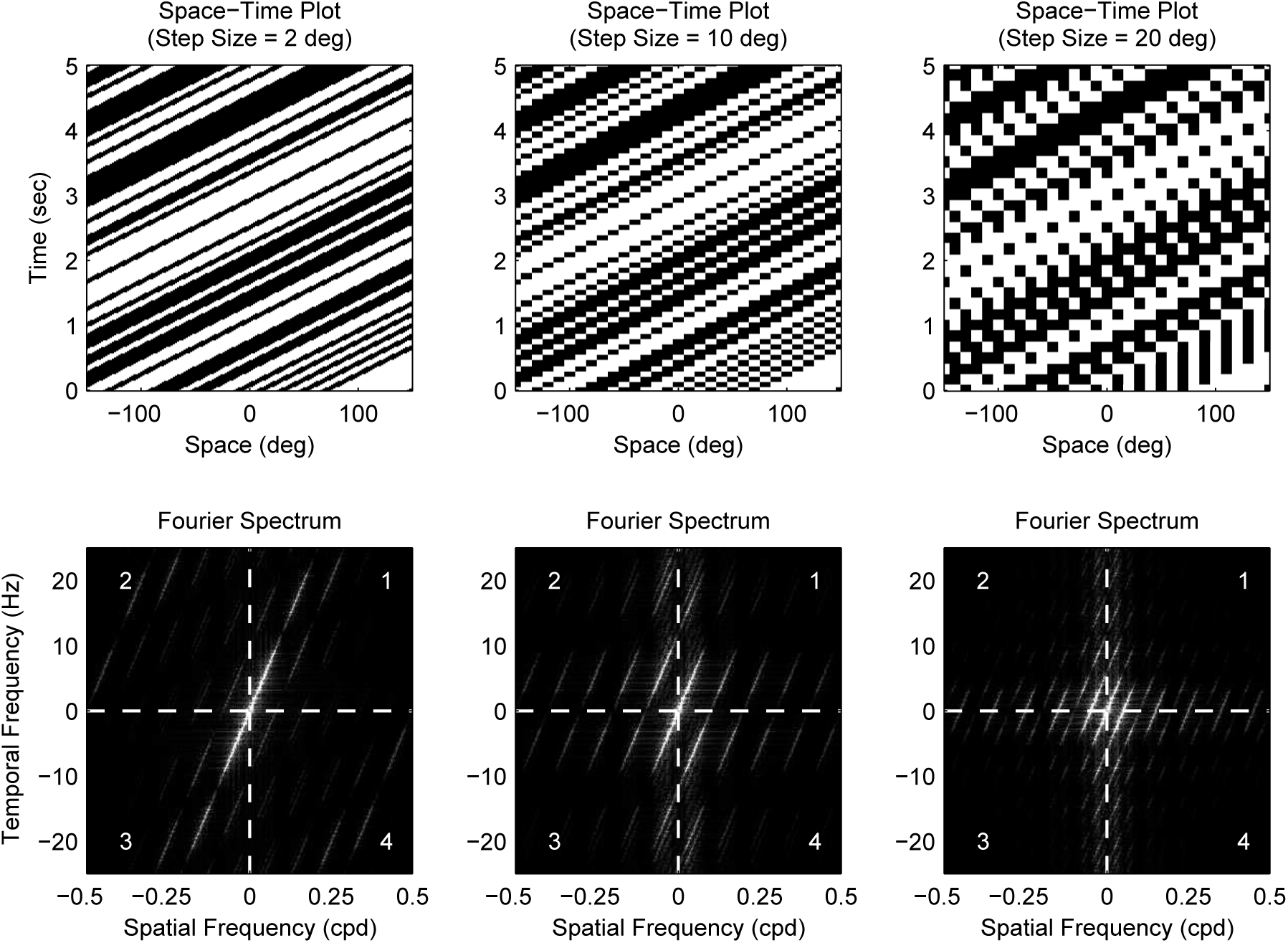
Stimuli with different step sizes (10 deg elements) Top panels show space-time plots and bottom panel corresponding spatiotemporal Fourier spectra for a number of step sizes (element size is 10 deg in all plots). When step size is much less than element size (left column), the stimulus is perceived by humans as moving smoothly and its motion energy is situated in quadrants 1 and 3 in the Fourier spectra (i.e. motion in the rightwards direction). In the middle column (step size equals element size), humans see the stimulus as moving less smoothly and part of its motion energy is now distributed in quadrants 2 and 4 in the Fourier domain (leftward motion). Finally, when step size is much larger than element size (right column), motion energy is more evenly distributed across the four Fourier quadrants and humans find it significantly more difficult to identify motion direction

An observer viewed the mantis through a web camera (while being blind to the stimulus) and coded its response in each trial as “moved left”, “moved right” or “did not move” (Nityananda et al. (2015)). Here, movement is defined as consistent optomotor response in the direction of the stimulus. This excludes trials in which mantids remained stationary or produced a peering response. The probability of motion was subsequently calculated as the proportion of trials in which motion was detected (mantis leaned in the same direction as the stimulus) under each condition. This is the closest feasible analogy to human psychophysics experiments, where humans classify their own perceptual experience into discrete classes. We have shown previously that human observers very rarely code the mantis as moving in the opposite direction to the stimulus Nityananda et al. (2015); Tarawneh et al. (2017). Consistent with our earlier work, we found that observers reported mantids to be moving in the opposite direction to the stimulus in only 4% of trials.

Figure 3 shows the collected data and fitted psychometric curves of motion probability versus step size for different element sizes. Comparing the different panels in Figure 3, it is clear that increasing element size shifts the psychometric curve in the positive direction of the step size axis. To quantify this change, we defined Dmax for each element size as the step size corresponding to a 50% probability of detecting motion, as estimated by a cumulative Gaussian psychometric function fit. Figure 4 shows pooled and individual plots of Dmax versus element size. We found that the relationship between Dmax and element size is in good agreement with the following power law (whose parameters we determined using maximum likelihood fitting):

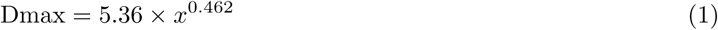
where *x* is element size in degrees.

**Figure 3:**
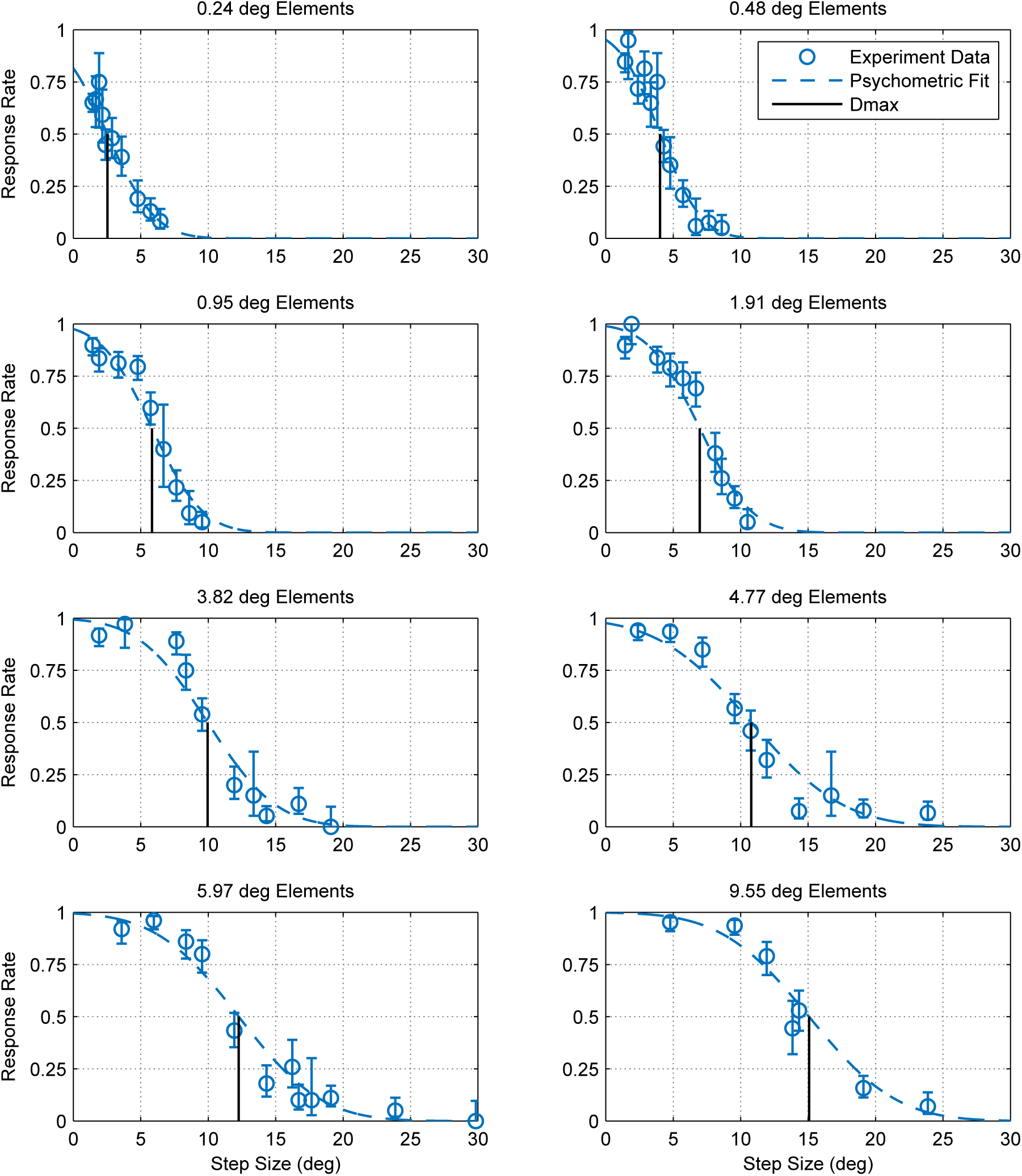
Optomotor response rates against step size for various element sizes. Error bars are 95% confidence intervals calculated using simple binomial statistics. Each panel shows the pooled responses of 13 mantids to a moving chequerboard stimulus of a given element size. The data and corresponding psychometric fits show that motion detection becomes increasingly more difficult (i.e. response rate decreases) as step size increases and that the step size corresponding to a 50% response rate (Dmax) increases with element size.

**Figure 4:**
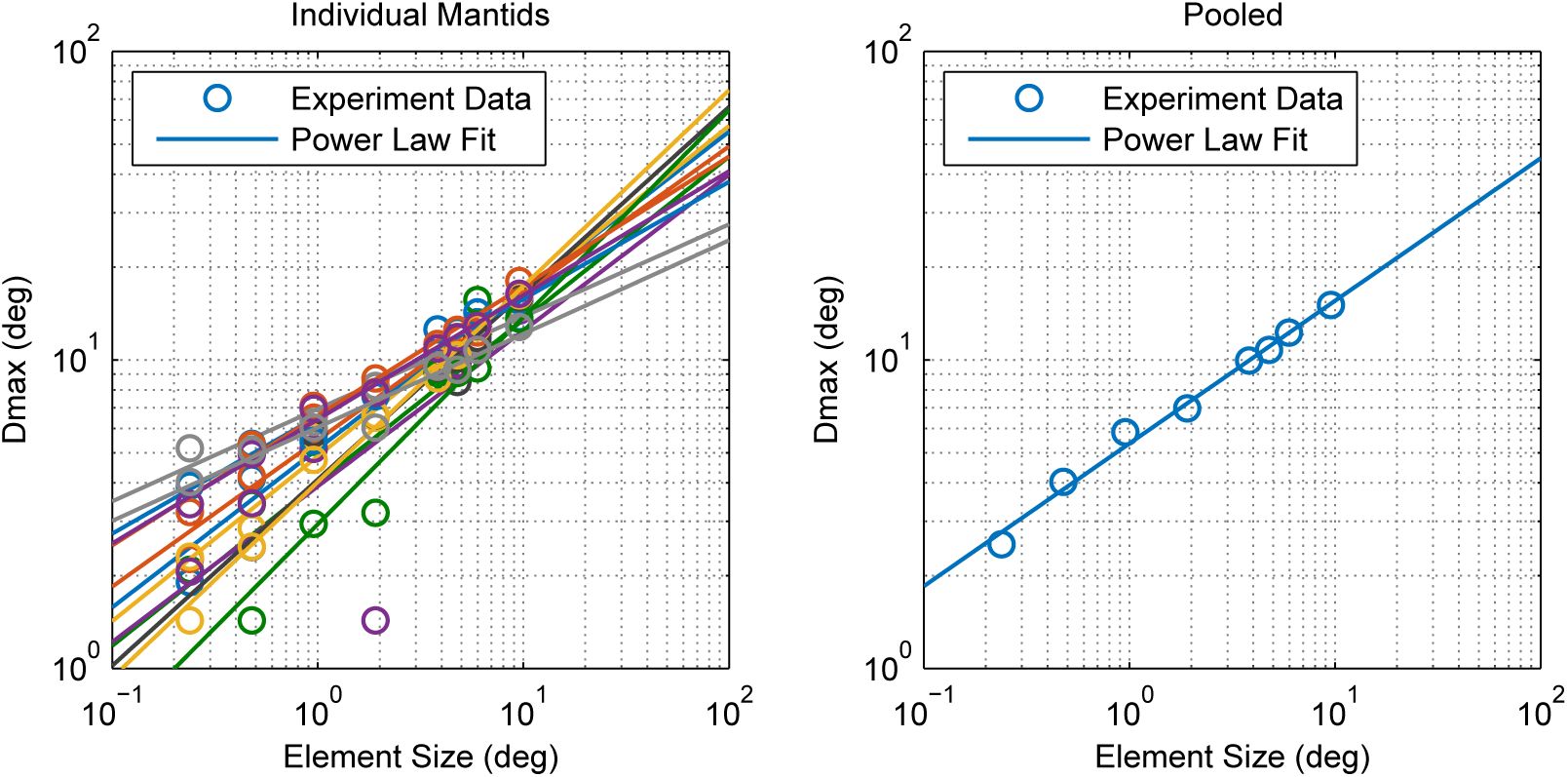
Relationship between Dmax and element size. Calculated Dmax values and and power law fits for individual mantids (left) and pooled data (right). The points appear to lie on a straight line on this log-log plot indicating that Dmax can be described accurately as a power law function of element size x (determined via fitting as Dmax(*x*) = 5.36 × *x*^0.462^ for pooled data).

### 2.2 Modeling and Simulation

#### 2.2.1 Simple Model based on Fourier Spectra

Modifying a pattern moving smoothly at a speed v by introducing a fixed step size ∆*x* is equivalent to passing it through a “sample and hold” circuit with a sampling interval of ∆*t* = ∆*x*/*v*. The transformation introduces an aliasing artifact whereby temporal frequencies higher than 1/2At are cast into lower frequencies as illustrated in Figure 2. A chequerboard pattern moving smoothly at a speed *v* consist of a series of components, each with a spatial frequency *f_s_* and a temporal frequency *f_t_* where *f_t_/f_s_* = *v*, that lie in quadrants 1 and 3 of the spatiotemporal Fourier domain (for motion in the positive direction). When the pattern is moved in discrete steps, a fraction of its component energy is transferred to quadrants 2 and 4 (i.e. motion in the negative direction). Increasing the step size ∆*x* (and correspondingly ∆*t*, since speed is constant) causes a larger portion of pattern motion energy to be distributed across the four quadrants, making it more difficult to identify the direction of motion. This aliasing effect can be observed in the space-time plots of moving patterns. When motion is smooth, the space-time plot shows clear rightward-pointings structures as in the left column of Figure 2. If the pattern is moved by a step size that is larger than element size (right column), leftward-pointing structures emerge as a result of the false matches between non-corresponding blocks across different frames.

As a first order approximation, the motion energy of a stepped chequerboard pattern can be taken as the sum of its unaliased components. The rationale behind this is that components in any temporal frequency range [*N −* 1/2, *N* + 1/2]/∆*t* (where *N* is a non-zero natural number) are cast by aliasing to the range [−1/2,1/2]/∆*t* and so, assuming symmetry around *N/*∆*t*, would be split into two groups with equal energy in the positive and negative directions. The net motion energy of aliased components in the pattern can therefore be approximated as zero (Figure 5). We ran numerical simulations in which we calculated the motion energy of stimuli in our experiment using this approximation. Figure 6 (left) shows a plot of approximated pattern energy versus the width of aliasing window ∆*f* (where ∆*f* = 1/∆*x*) for a number of element sizes. The plot shows that increasing element size causes a proportional shift in the curve over a region of energy levels (up to ~ 2 × 10^5^ units). Assuming the energy threshold T corresponding to Dmax is within this range (e.g. *T* ~ 10^5^), we calculated Dmax as the step size at which motion energy crosses the threshold (for each element size). Figure 6 (right) shows a plot of calculated Dmax versus element size based on this model and compares this with our experimental findings. The predictions and experimental results are in good agreement, showing that this simple model with a single parameter, the threshold *T*, can account well for the Dmax values we observed in the mantis.

**Figure 5:**
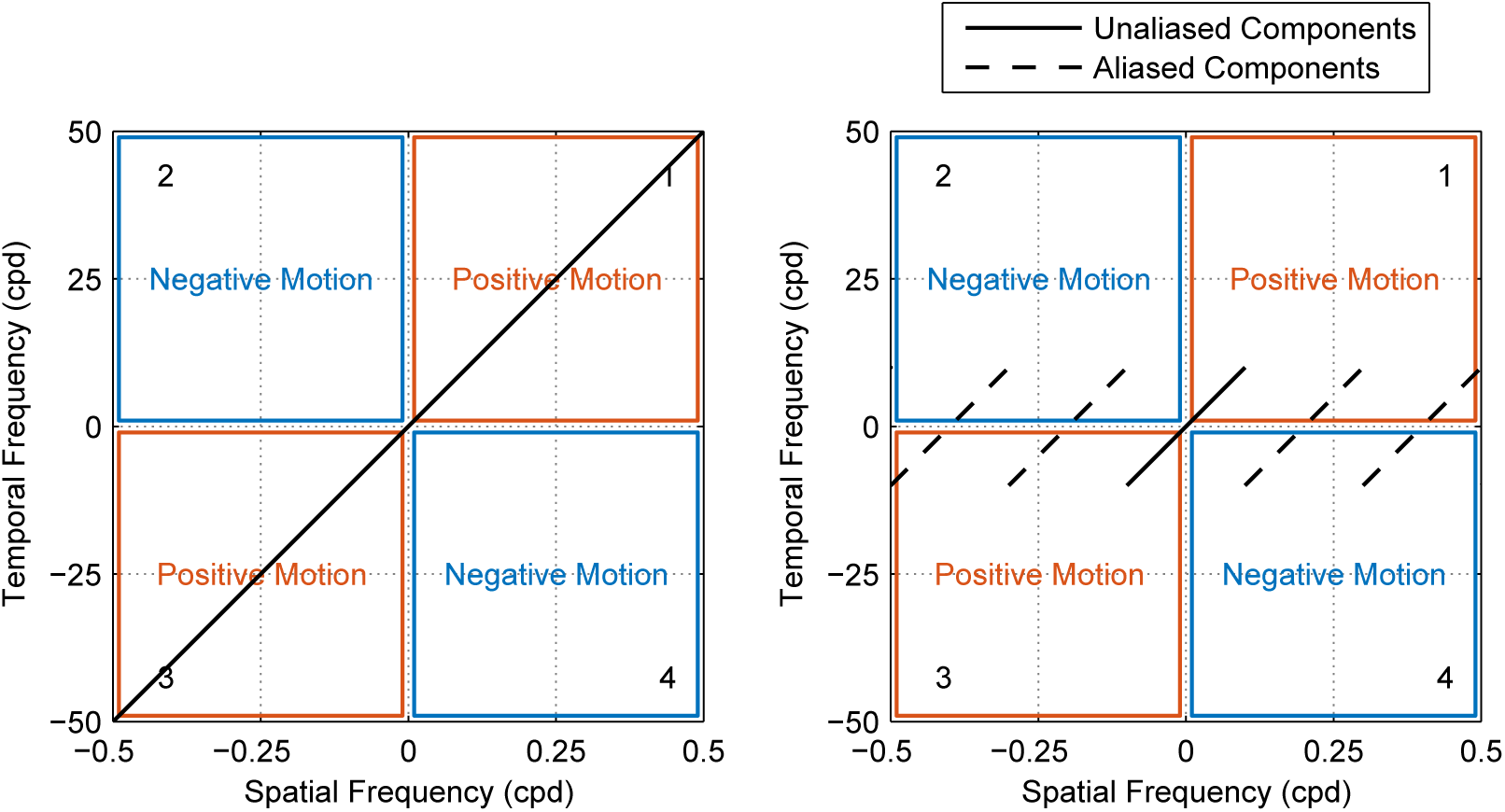
Effect of aliasing on the distribution of motion energy in the Fourier domain. A pattern moving coherently at a constant speed v has spatial and temporal frequency components along the line *f_t_/f_s_* = *v* and its motion energy content is in quadrants 1 and 3 that signify positive motion (left panel). When the pattern is moved in steps of ∆*x* at the same speed, components with temporal frequencies higher than *v*/∆*x* get aliased and cast towards lower frequencies (right panel). Aliased components are distributed in the four quadrants and their net motion energy is therefore close to zero, leaving the energy of unaliased components as a close approximation of what remains in quadrants 1 and 3.

**Figure 6:**
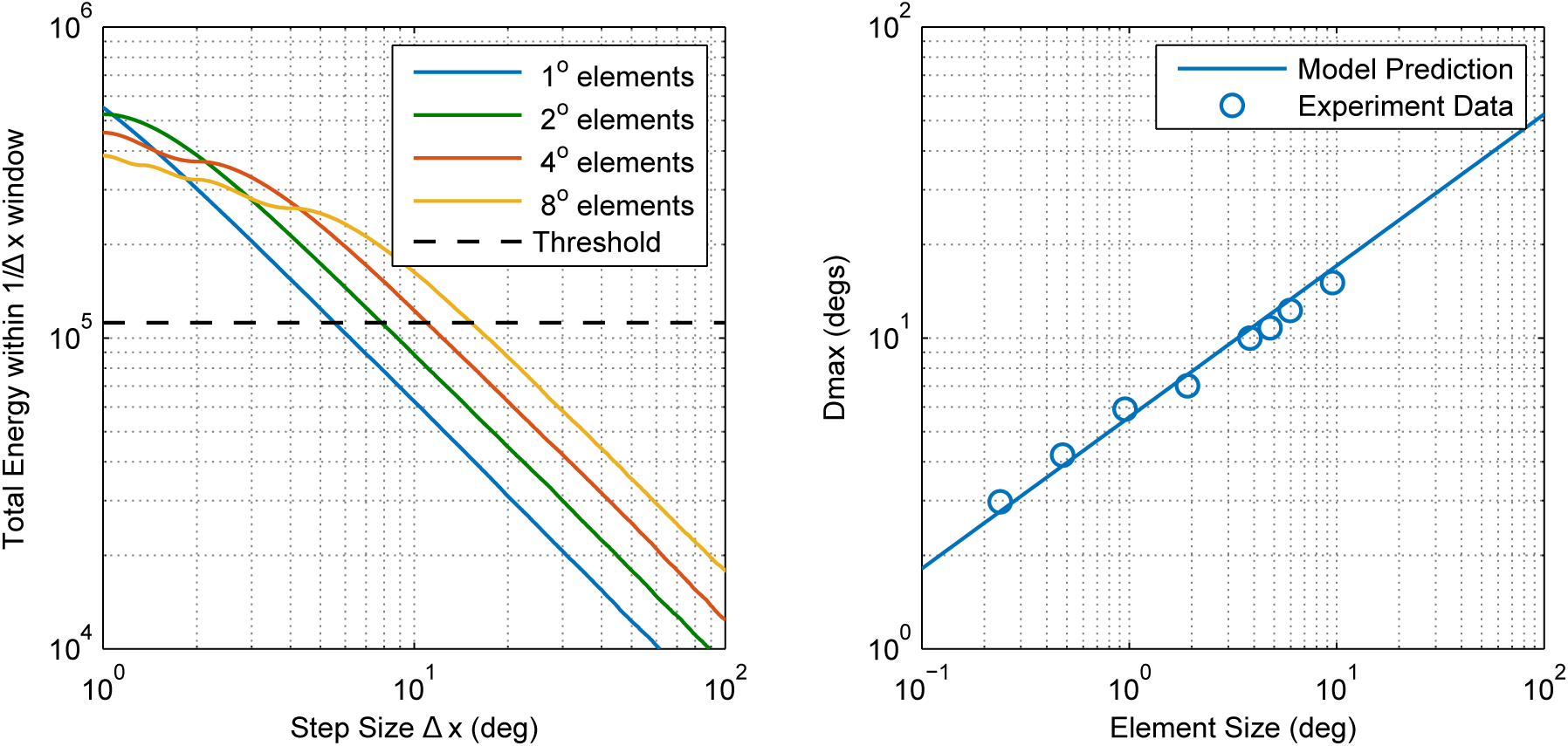
Simple motion energy-based model of Dmax in the praying mantis. (Left) the total motion energy of unaliased pattern components contained within a frequency window [−1/∆*x*, 1/∆*x*] decreases as the step size ∆*x* is increased. Patterns of larger elements have a larger portion of their motion energy within lower frequencies and therefore contain more energy at a given step size. According to this model, Dmax is the step size corresponding to a given motion energy threshold (chosen here as ~ 10^5^ units). (Right) This simple model is in good agreement with our experimental results for the mantis.

#### 2.2.2 Reichardt Model

Our simple model based on the Fourier spectra of stimuli provides a good quantitative account of Dmax but assumes an underlying motion detection system that is equally sensitive at all spatial and temporal frequencies. Published data of the mantis contrast sensitivity function, however, show maximum sensitivity in the spatial frequency range 0.01 to 0.1 cpd (Nityananda et al., 2015) which is at odds with what our simplified model based on Fourier energy assumes. We therefore built and simulated a lower-level motion detection model to investigate whether our experimental findings could be accounted for by a system of narrow-band detectors. This second model is based on the well known Reichardt Detector (RD) that is widely used to account for the optomotor response and similar large-field visually-guided forms of behavior in insects (Bahl et al., 2013; Borst and Bahde, 1988; Hassenstein and Reichardt, 1956; Srinivasan et al., 1991). The detector (shown in Figure 7) consists of two subunits that compute motion in opposite directions, typically assumed to be neighboring ommatidia and their neural cartridges (Borst et al., 2010). Spatial input from each subunit is temporally delayed then compared with the other subunit’s input to detect luminance changes that have “travelled” in the same direction with a similar delay. The two subunit outputs are then summed to produce a direction-sensitive measure of motion.

**Figure 7:**
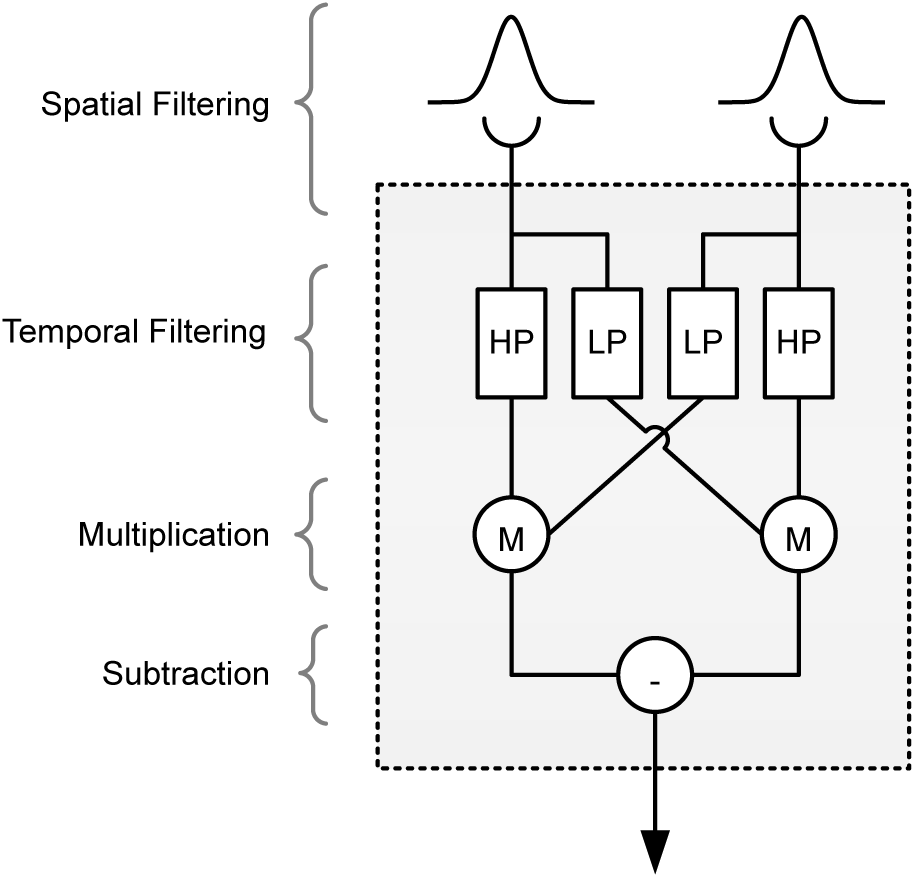
The Reichardt Detector

Although the preferred spatial frequency of the detector can be varied by adjusting the spatial separation of its subunits and the extent of their input spatial filters, its spatial bandwidth is considerably constrained (Tarawneh et al., 2017). To enable the model to respond to stimuli covering a spatial bandwidth larger than the detector’s (such as the range of element sizes we tested), we therefore combined multiple detectors tuned to two spatial frequencies (i.e. two detector *classes*). Our model is shown in Figure 8 and is essentially a slight variation of a published model of the mantis optomotor response (Tarawneh et al., 2017). The (free) parameters that we fitted were the separation and extent of spatial filters for each class (*σ*_1_, *σ*_2_, ∆*x*_1_, ∆*x*_2_), the two post-detector thresholds *T*_1_ and *T*_2_ (each for one of the two classes) and the magnitude of mantis and observer noise levels. The values of other parameters including temporal tuning characteristics of the RD were selected based on physiologically plausible data reported in literature (Borst (2014b); Nityananda et al. (2015); Tarawneh et al. (2017)).

**Figure 8:**
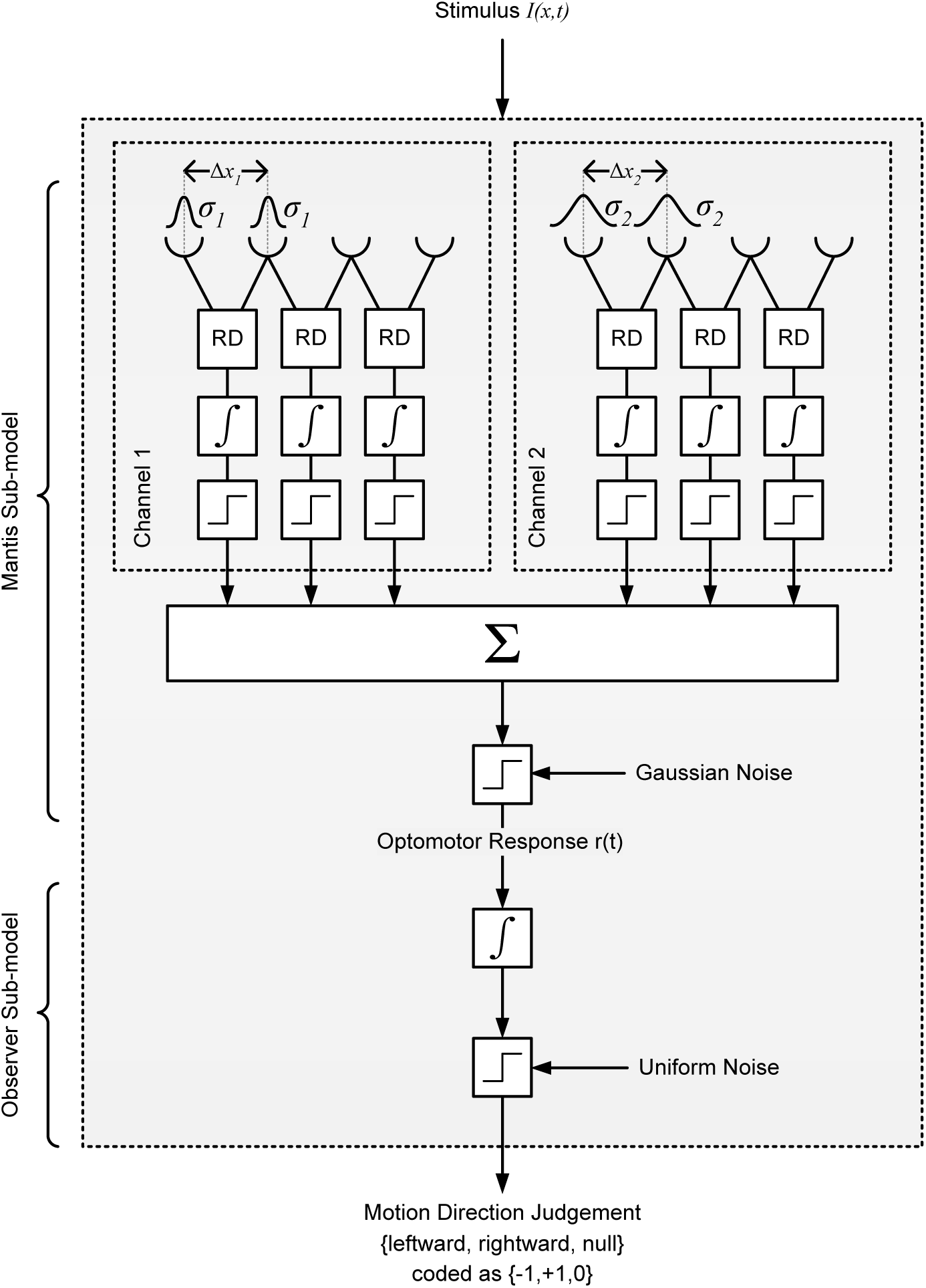
Mantis optomotor response model based on the Reichardt detector. The model has two arrays of detectors that are positioned in random locations across a simulated 1D retina. Detectors within each array share the same subunit separation and spatial filter extent (and are thus all tuned to the same spatial frequency). Having two detector classes enables to model to respond to stimuli across a range of spatial frequencies broader than that of a single detector. The output of each detector is temporally integrated and then converted to a “vote” for leftward, rightward or no motion by a hard threshold. The votes are subsequently combined, thresholded and then passed through another temporal integrator and hard threshold blocks that model the decision process of the human observer in the experiment

Figure 9 shows the predictions of this model (obtained by running simulated trials using the same stimuli as the experiment) versus experiment data. The predictions are in good agreement with our experimental observations showing that, by combining at least two classes of detectors, mantis Dmax values can be explained by a system of narrow-band motion detection units.

**Figure 9:**
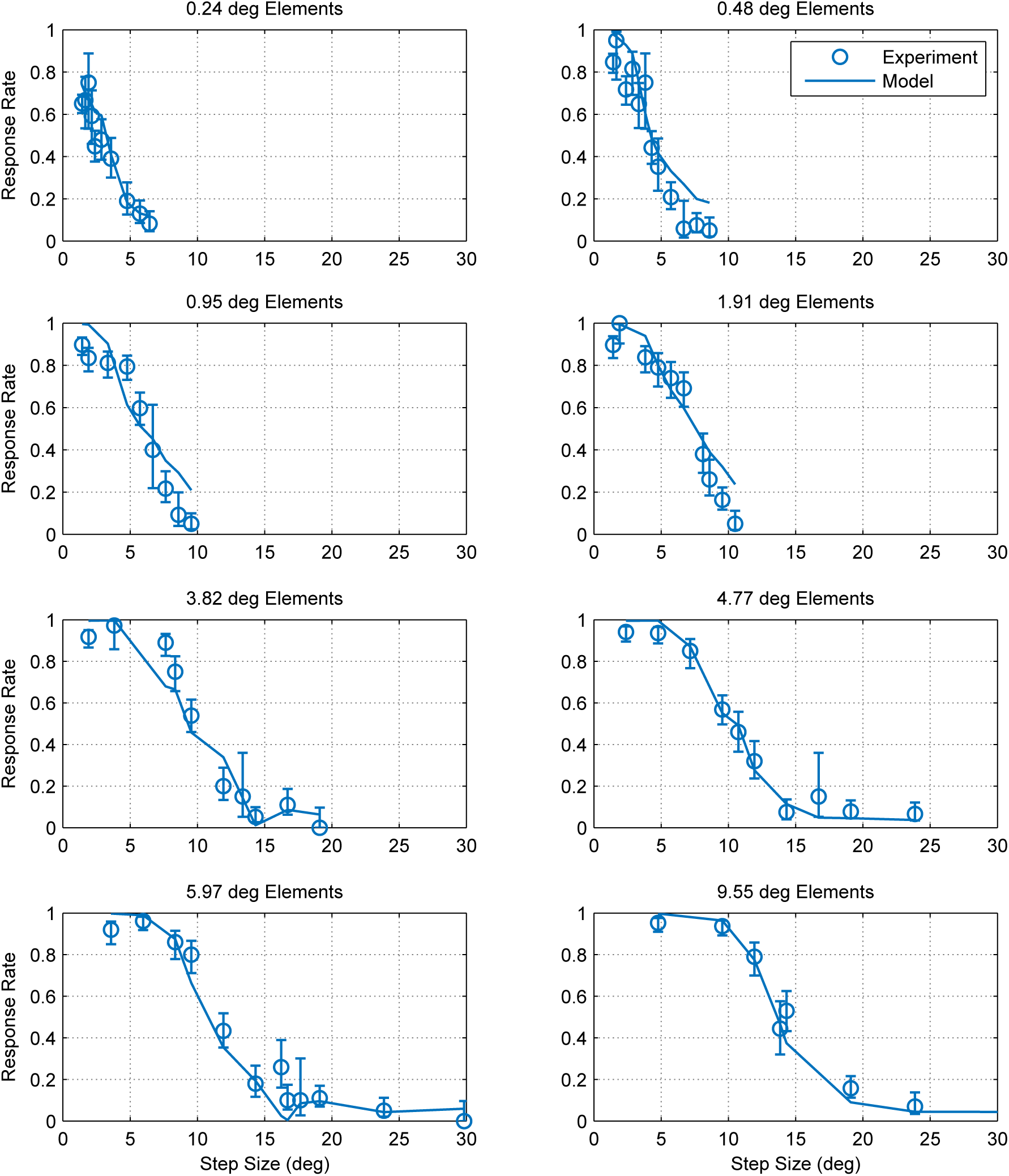
Predictions of the RD-based model against experimental results

## 3 Discussion

Our results show similarities and differences between Dmax in humans and mantids. First, while mantis Dmax appears to scale with element size similar to humans (for elements larger than 15 arcmin), we did not observe a region of element sizes with a constant Dmax (up to 15 arcmin in humans). Humans can perceive static dot patterns with much smaller element sizes and so the lower bound on Dmax is taken as evidence for the existence of a spatial filtration stage that precedes motion detection (Morgan, 1992). Consistent with this, applying spatial low pass filtration to random dot stimuli was found to increase the lowest Dmax bound (Morgan, 1992). Although it is possible that element sizes smaller than the ones we tested may have a constant Dmax, the smallest element size in our experiment (0.24 deg) was smaller than the foveal inter-ommatidial angle in the comparable mantis species *Tenodera australasiae* (Rossel, 1979). Mantids are also most sensitive to motion with spatial frequencies of 0.01 ~ 0.1 cpd (Nityananda et al., 2015), a range much lower than the fundamental spatial frequency component of a chequerboard pattern with 1 deg elements (2 cpd). Patterns with elements as small can still be detected because of their lower frequency components but, as element size decreases further, a larger portion of their energy shifts towards the highest spatial frequencies and outside the mantis sensitivity range. It is therefore likely that the smallest element size we tested is near the limit of mantis visual acuity. Consistent with this, our data (Figure 3A) show that the response rate for 0.24 deg elements is significantly lower, even at the smallest step size, compared to larger elements. Mantids may therefore have no range of element sizes over which Dmax is constant.

Another apparent difference between mantis and human data concerns the relationship between Dmax and element size. In humans, Dmax is assumed to scale linearly with element size (i.e. Dmax = *kx*) (Cavanagh et al., 1985; Morgan, 1992; Smith and Ledgeway, 2001; Watanabe, 1998) where the coefficient of proportionality *k* depends on a number of stimulus parameters such as spatial extent (Baker and Braddick, 1982; Chang and Julesz, 1983; Nakayama and Silverman, 1984; Todd and Norman, 1995) and duration of presentation (Nakayama and Silverman, 1984; Nishida and Sato, 1992; Snowden and Braddick, 1989a,b). Our results in the mantis however appear to be best approximated by a power law fit with an exponent of of 0.462 (i.e. Dmax = *kx*^0.462^). Even though the coefficient of proportionality *k* is also likely to depend on stimulus parameters, the different exponent may reflect an actual difference between motion processing in humans and mantids. However, a comparison between mantis and published human Dmax data appears to show that this difference is not very pronounced (Figure 10). Fitting *k* to all the human data while constraining the exponent to be 0.462, as for mantids, produces a reasonable fit, even neglecting the lower bound on Dmax at small element sizes. In a more complex model accounting for this limit, it seems far from clear that an exponent of 1 would be the best fit. In fact, Dmax in both humans and insects may show a sublinear dependence on scale.

**Figure 10:**
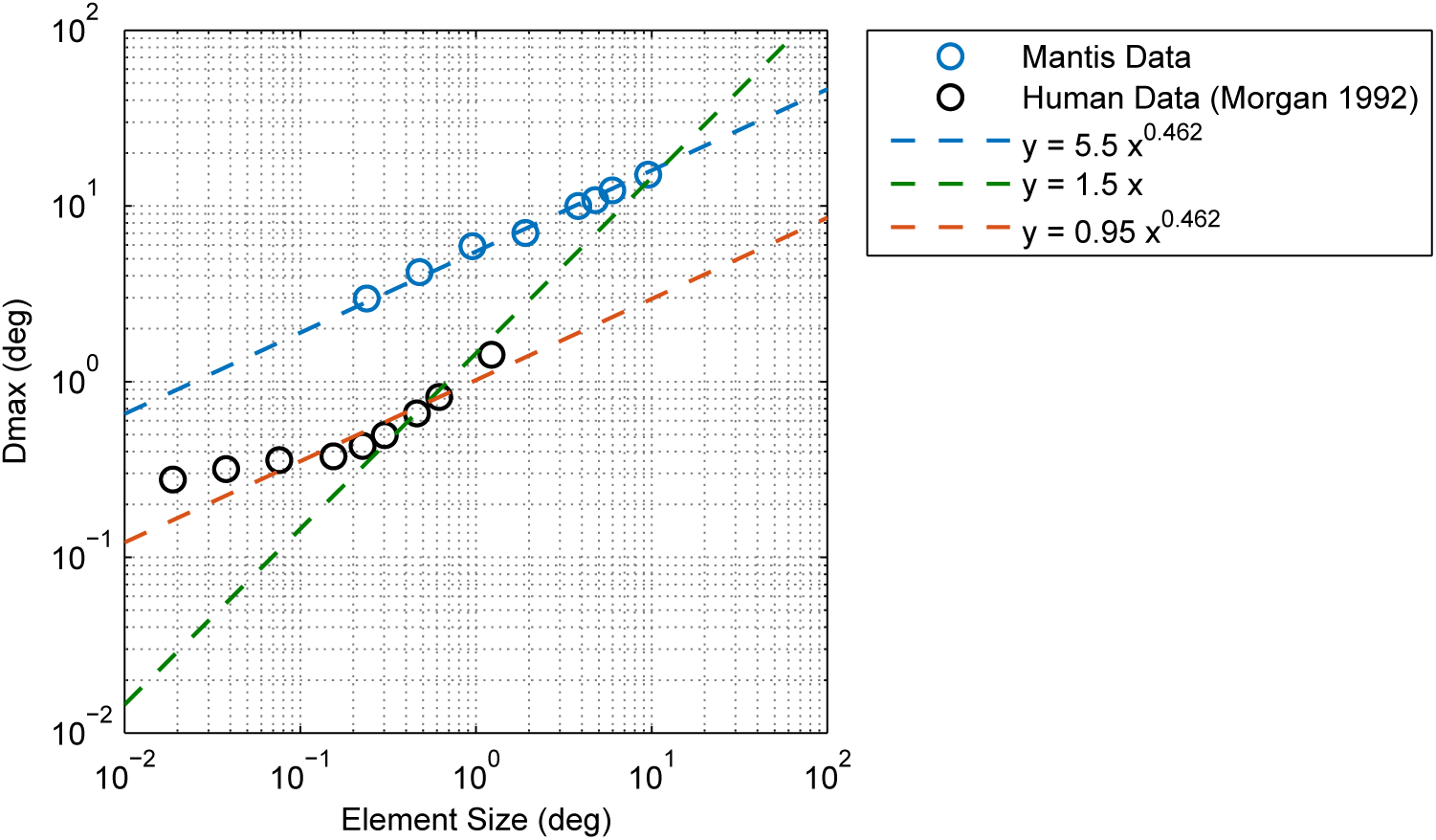
Comparison between Dmax in humans and mantids. Mantis Dmax increases with element size similar to humans but does not appear to have a lower limit below which it remains constant (in humans, this limit is about 15 arcmin = 0.25 deg). The relationship between mantis Dmax and element size is well accounted for by a power law function with an exponent of 0.462 (blue line). Human data from Morgan (1992) is plotted for comparison. Morgan interprets the Dmax values he observed as scaling proportionately for element sizes larger than 0.25 deg (i.e. assumes a power law exponent of 1, green line) although his data appears to be accounted for equally well by a power law with an exponent of 0.462 (red line), similar to mantids. It is therefore unclear whether a true difference in power law exponents exists between humans and mantids.

The ability of both our Fourier and RD-based models to reproduce the relationship between Dmax and element size in the mantis suggests further similarities. Our simple model based on stimulus energy in the Fourier domain is similar to the Motion Energy model used to account for apparent motion perception in humans (Adelson and Bergen, 1985). Both assume that motion perception is computed by a cross-correlation operation in the space-time domain (or, equivalently, a summation of stimulus energy in the spatio-temporal Fourier domain). A direct implication of our model is that Dmax corresponds to a motion energy detection threshold that is reliably exceeded by smoothly moving stimuli but not by those with aliasing artifacts caused by large motion steps. According to this model, Dmax scales with element size because stimuli composed of larger elements have a greater portion of their energy within lower frequencies and can therefore be more severely aliased by larger step sizes before their unaliased component energy falls below threshold. This simple account of Dmax is sufficient to explain the main characteristics of the relationship between Dmax and element size but does not take into account the limited spatial sensitivity of local correlation-type detectors. The response properties predicted by these detectors were found to be in excellent agreement with a series of behavioral and physiological studies (Eckert, 1973; Joesch et al., 2008, 2010; Rister et al., 2007; Schnell et al., 2010; Von Fermi and Richardt, 1963) and much is known about their neural substrates (Borst, 2014b). Reproducing the relationship between Dmax and element size using the RD-based model was therefore important to verify that the intuition behind the Fourier model still holds when motion detection is mediated by units with narrow-band sensitivity and other physiologically-realistic details.

In conclusion, even though we observed that mantis Dmax scales non-linearly with element size *x* (specifically that Dmax = *x*^0^·^462^) and that it does not appear to have a minimum bound, these observations appear consistent with published human Dmax data nonetheless. We also found that two models based on the RD and motion energy (that are predominantly used in insect and human motion detection, respectively) provided consistent accounts of these experimental observations. Our findings add to the body of evidence suggesting that similar computational mechanisms underlie visual motion detection in insects and mammals (Adelson and Bergen (1985); Borst and Helmstaedter (2015); Lu and Sperling (1995); Tarawneh et al. (2017); Van Santen and Sperling (1984)).

## 4 Methods

### 4.1 Insects

A total of 13 female mantids of the species *Sphodromantis lineola* were used in the experiment. Each individual was kept in a plastic box of dimensions 17 × 17 × 19 cm with a porous lid for ventilation and fed a live cricket twice per week. The boxes were kept at a temperature of 25° and were cleaned and misted with water twice per week.

### 4.2 Experimental Setup

In each experiment, a mantis hung upside down from a Perspex base and viewed full-screen stimuli on a CRT monitor (HP P1130). The base was held in place by a clamp such that the mantis was 7 cm away and facing the middle point of the screen. A web camera (Kinobo USB B3 HD Webcam) was placed under the base providing a view of the mantis (but not the screen). The monitor, Perspex base and camera were all placed inside a wooden enclosure to isolate the mantis from distractions and maintain consistent dark ambient lighting during experiments. The screen had physical dimensions of 40.4 × 30.2 cm and pixel dimensions of 1600 × 1200 pixels. At the viewing distance of the mantis the horizontal extent of the monitor subtended a visual angle of 142°. The mean luminance of the monitor was 27.1 cd/m^2^ and its refresh rate was 85 Hz.

Experiments were programmed in Matlab 2012b (Mathworks, Inc., Massachusetts, US) and used stimuli developed using Psychophysics Toolbox Version 3 (PTB-3) (Brainard and Vision, 1997; Kleiner et al., 2007; Pelli, 1997). The experiment computer was a Dell Optiplex 9010 (Dell, US) with an Nvidia Quadroo K600 graphics card running Microsoft Windows 7.

### 4.3 Experimental Procedure

Each experiment consisted of a number of randomly-interleaved trials in which a mantis was presented with moving chequerboard patterns. An experimenter observed the mantis through the camera underneath and blindly coded the direction of the elicited optomotor response (if any). The response code for each trial was “moved left”, “moved right” or “did not move”. There were equal repeats of left-moving and right-moving patterns of each condition in all experiments. At the beginning of each trial, an alignment stimulus was used to steer the mantis back such that it is facing the middle point of the screen. The alignment stimulus was a smoothly moving chequerboard pattern controlled by the experimenter using a keyboard.

### 4.4 Visual Stimulus

The stimulus was a full-screen chequerboard pattern moving either leftwards or rightwards in each trial. The pattern was composed of black and white chequers of 0.161 and 54.1 cd/m^2^ luminance levels respectively, chosen randomly with equal probability. The whole pattern was displaced horizontally by a step ∆*x* at ∆*t* intervals where ∆*x*/∆*t* was kept constant across the different conditions.

### 4.5 Calculating Dmax

We used likelihood fitting to calculate Dmax as the step size corresponding to a 50% response rate for each element size. The fitted psychometric functions were in the form:

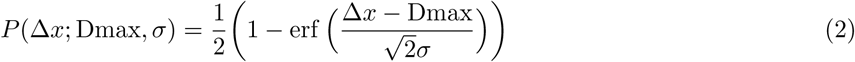
where *p* is the probability that motion is detected, ∆*x* is step size and *σ* is a parameter that specifies the width of the function’s transition period. The parameters Dmax and *σ* were calculated for each element size using maximum likelihood estimation in Matlab, assuming mantids’ responses had a simple binomial distribution. Therefore:

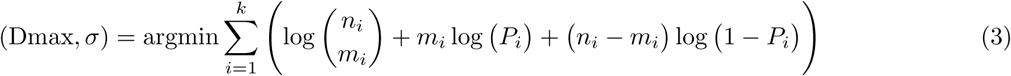
where *k* is the number of step sizes, *n_i_* is the number of trials in which mantids detected motion, *m_i_* is the total number of trials and *P_i_* is motion detection probability predicted by given Dmax and *σ* values as per Equation 2. We calculated Dmax and *σ* for each insect’s individual data and for all insects pooled together.

## 5 Data Availability

The data collected in this study are available on https://github.com/m3project/mantis-dmax

